# Enterohemorrhagic Escherichia coli (EHEC) Disrupts Intestinal Barrier Integrity in Translational Canine Stem Cell-Derived Monolayers

**DOI:** 10.1101/2024.02.27.582360

**Authors:** Itsuma Nagao, Minae Kawasaki, Takashi Goyama, Hyun Jung Kim, Douglas R. Call, Yoko M. Ambrosini

## Abstract

This study addresses the gap in translatable *in vitro* models for investigating Enterohemorrhagic *E. coli* (EHEC) infections, particularly relevant to both canine and human health. EHEC is known to induce acute colitis in dogs, leading to symptoms like hemorrhagic diarrhea and hemolytic uremic syndrome, similar to those observed in humans. However, understanding the pathophysiology and developing treatment strategies have been challenging due to the lack of effective models that replicate the clinical disease caused by EHEC in both species. Our approach involved the development of colonoid-derived monolayers using intestinal tissues from healthy, client-owned dogs. These monolayers were exposed to EHEC and the impact of EHEC was assessed through several techniques, including trans-epithelial electrical resistance (TEER) measurement, immunofluorescence staining for junction proteins and mucus, and scanning electron microscopy for morphological analysis. Modified culture with saline, which was intended to prevent bacterial overgrowth, maintained barrier integrity and cell differentiation. EHEC infection led to significant decreases in TEER and ZO-1 expression, but not in E-cadherin levels or mucus production. Additionally, EHEC elicited a notable increase in TNF-α production, highlighting its distinct impact on canine intestinal epithelial cells compared to non-pathogenic *E. coli*. These findings closely replicate *in vivo* observations in dogs and humans with EHEC enteropathy, validating the canine colonoid-derived monolayer system as a translational model to study host-pathogen interactions in EHEC and potentially other clinically significant enteric pathogens.

## Introduction

Human acute colitis is frequently attributed to food-borne infections caused by Enterohemorrhagic *Escherichia coli* (EHEC) that produce Shiga toxin (Stx). This toxin induces cytotoxic effects on adjacent endothelial cells, resulting in symptoms like bloody diarrhea and hemolytic uremic syndrome (HUS) (1). While studying EHEC, researchers have encountered challenges in modeling the disease. Mouse models, for example, failed to replicate the expected effects of mucosal colonization and intestinal damage (2).

Although the pathogenicity of EHEC in dogs are relatively mild (3), acute colitis due to EHEC infection, including hemorrhagic diarrhea and HUS similar to humans, has been demonstrated in experimentally infected dogs or critically ill dogs (4, 5). Because of this similarity, there is a growing interest in using dogs as *in vivo* models for human EHEC infections and as sources of cells and tissues for *in vitro* models.

Recent advancement of organoid technology offers improved *in vitro* models compared to traditional cell culture systems (6, 7). Organoid models include three-dimensional intestinal organoids. Organoids derived from adult stem cells closely mimic the *in vivo* microenvironment and maintain multi-cell lineage cultures (7). Two-dimensional (2D) monolayers can also be derived from organoids (8). These 2D monolayers offer a stable and accessible luminal interface to exam nutritional effects, drug permeability, and host-pathogen interactions. Additionally, sophisticated microfluidic intestine-on-a-chip systems have been established to study changes in the intestinal microenvironment and host responses with more physiologically complex *in vitro* models (9, 10).

While there is a wealth of literature on murine and human intestinal organoids and associated technologies, research on canine equivalents is still emerging (11). Canine models are gaining prominence due to their relevance as a spontaneous, large animal model for various human intestinal diseases, including EHEC infection. Therefore, the present study aimed to establish a comparative *in vitro* canine 2D monolayer model to assess the effects of EHEC infection, utilizing canine colonoid-derived monolayers and a strain of non-pathogenic *E. coli* as an infection control.

## Results

### Multilineage cell differentiation and intestinal barrier integrity of monolayers were maintained with saline

In this study, we modified the apical medium 24 hours before infection by switching from the standard organoid culture medium to a nutrient-free saline solution. This adjustment aimed to prevent bacterial overgrowth in the medium, as previously reported (12) (Figure 1A). Initially, we assessed whether the colonoid-derived monolayers, which had been cultured in the organoid culture medium for four days, could be sustained in saline for three days, aligning with the desired duration of co-culture with bacteria. During the three-day saline culture period, the colonoid-derived monolayers maintained their confluency (Figure 1B), and formed functional tight junctions (Figure 1C). To further confirm the presence of differentiated goblet cells, we conducted staining using Sambucus Nigra Agglutinin (SNA), which is a lectin that identifies sialic acid bound to galactose in mucin (13). This staining revealed the presence of SNA-positive cells on the colonoid-derived monolayers (Figure 1C). Additionally, to verify the multi-lineage characteristics of the monolayers, we utilized quantitative polymerase chain reaction (qPCR). This technique was instrumental in determining whether there was a variation in the composition of differentiated cells between two groups: one being colonoid-derived monolayers cultured in nutrient-free saline, and the other in standard organoid culture medium. Our analysis revealed no significant differences between the two groups for the expression levels of stem cell marker *leucine-rich repeat containing G protein-coupled receptor 5* (*LGR5*), the enteroendocrine cell marker *Chromogranin A*, and the Paneth cell marker *Lysozyme* (Supplementary Figure 1).

**Figure 1.**
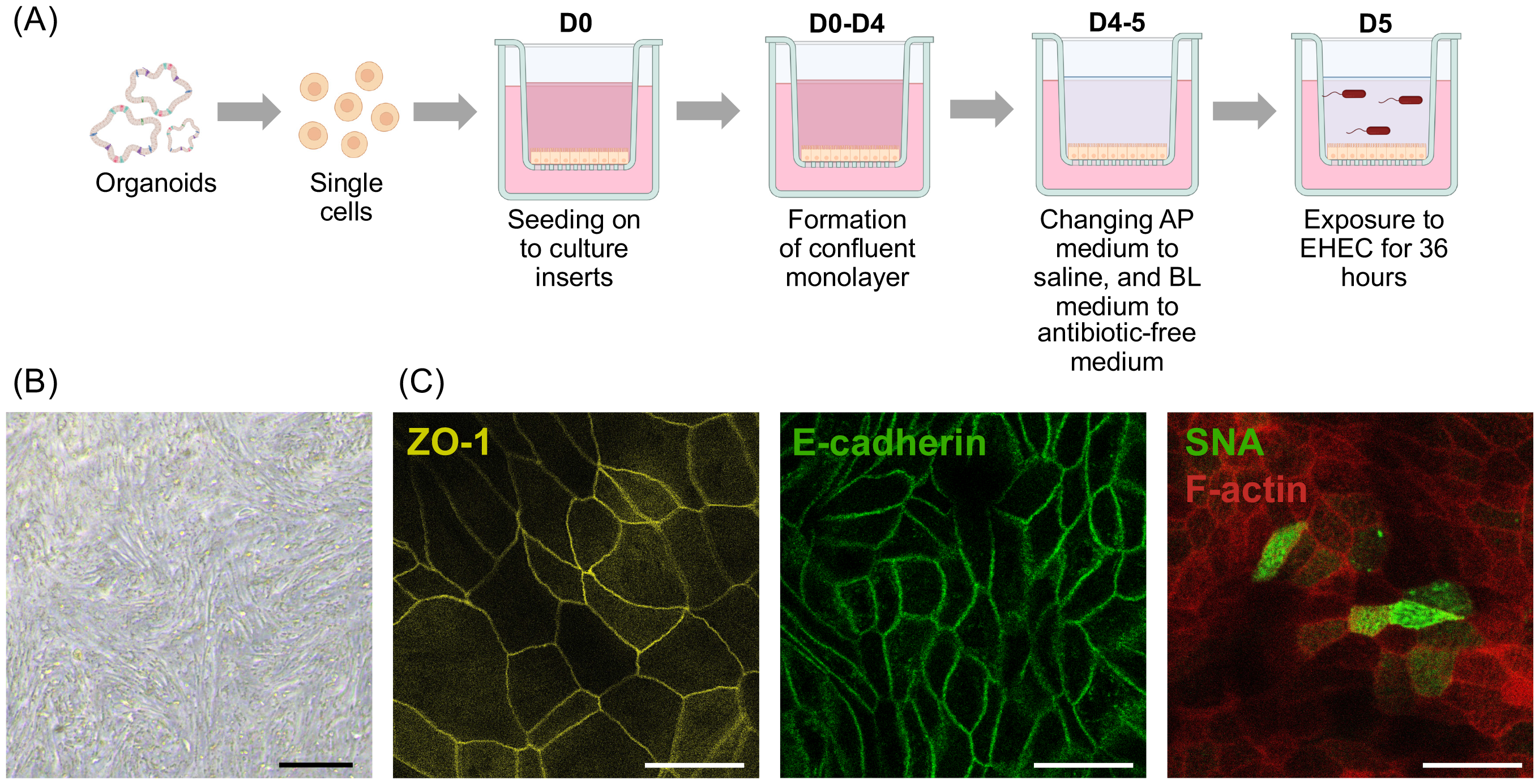
Modification of canine colonoid-derived monolayers for investigating EHEC infection. (A) A schematic diagram of the experiment. Colonoid-derived monolayers were derived from canine colonoids (D0-D4), and organoid culture medium was replaced with saline in apical chamber to avoid over-growth of bacteria and antimicrobial-free organoid culture medium in basolateral chamber once the monolayers reached stable state at day 4 (D4). Then, co-culture with EHEC was conducted on day 5 (D5). This schematic was created with BioRender.com. (B) Representative phase-contrast image of colonoid-derived monolayers after 3 days culture using saline. Scale bar = 100 μm. (C) Representative fluorescence image of colonoid-derived monolayers after 3 days culture using saline. Immunofluorescence staining on canine colonoid-derived monolayers confirms expression of tight junction protein, ZO-1 (Yellow) and E-cadherin (Green). The population of goblet cells (SNA: Green) was also visualized by using immunofluorescence staining with a counterstaining, F-actin (Red. Scale bar = 25 μm.

### Establishment of EHEC co-culture on canine colonoid-derived monolayers

Next, EHEC isolated from cattle was co-cultured with the colonoid-derived monolayers. Because clinical isolates of EHEC from dogs were not available, a clinical isolates from cattle was used. In addition, a non-pathogenic *E. coli* strain, MC4100, was used as an infection control strain for *E. coli* infection (14). After 36 hours of infection, microvilli extended uniformly across the surface of the epithelium in all uninfected control cultures, and in cultures infected with non-pathogenic *E. coli* or EHEC (Figure 2). Bacteria were adhering to the cell surface for both non-pathogenic *E. coli* and EHEC cultures, but the attachment of bacteria to the cell surface was different. Non-pathogenic *E. coli* adhered to the cell surface but did not bind to the microvilli, while EHEC was observed to gather surrounding microvilli and bind strongly to the cell surface (Figure 2).

**Figure 2.**
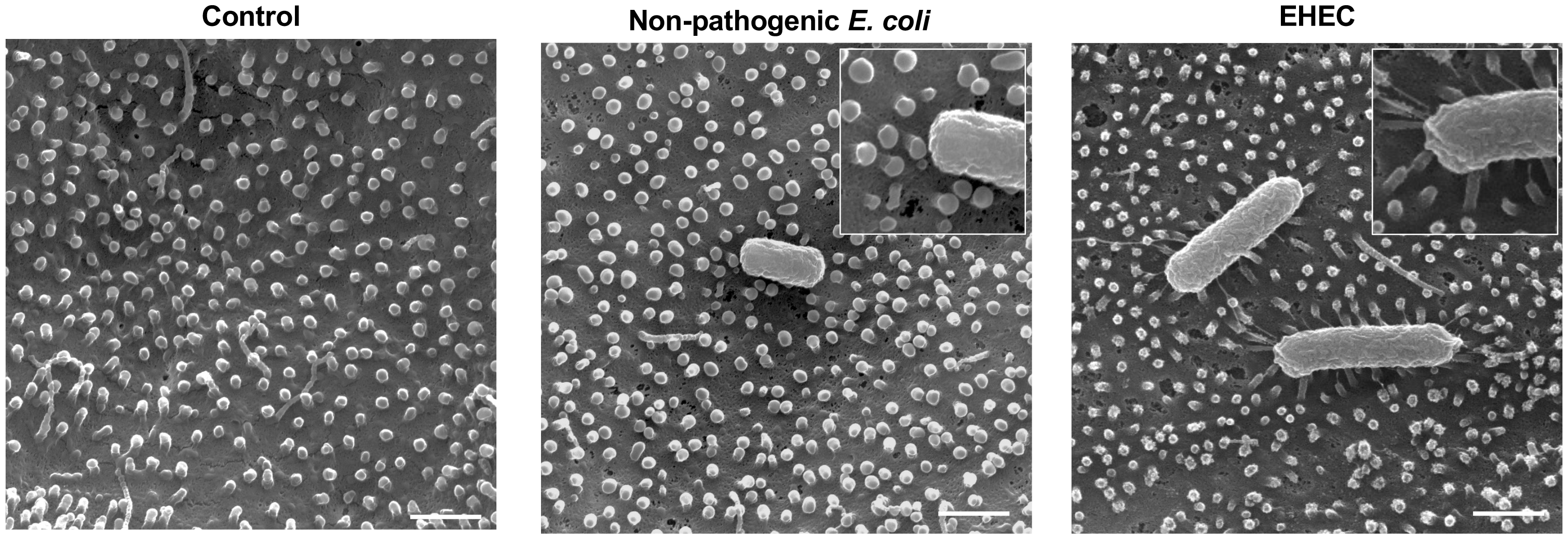
Scanning electron microscopy images of canine colonoid-derived monolayers co-cultured with bacteria. After 36 hours of co-culture, the cells were fixed and their surfaces were observed with a scanning electron microscope. Representative images of uninfected control, non-pathogenic *E. coli*, and EHEC are shown on the left, middle, and right, respectively. Scale bar = 1 μm.

Phase-contrast microscopy indicated no major differences in appearance between cultures treated with non-pathogenic *E. coli* or EHEC, with both maintaining confluent monolayers (Figure 3A). Trans-epithelial electrical resistance (TEER) values decreased from the start of culture for all cultures for the first 24 hours but were maintained in the uninfected control and non-pathogenic *E. coli* cultures thereafter. In contrast, the TEER values for the EHEC infected monolayers continued to decline, showing a significant decrease compared to the non-pathogenic *E. coli* infection at 32 hours and 36 hours post-coculture (Figure 3B) (*P*=0.03 and *P*=0.01, respectively).

**Figure 3.**
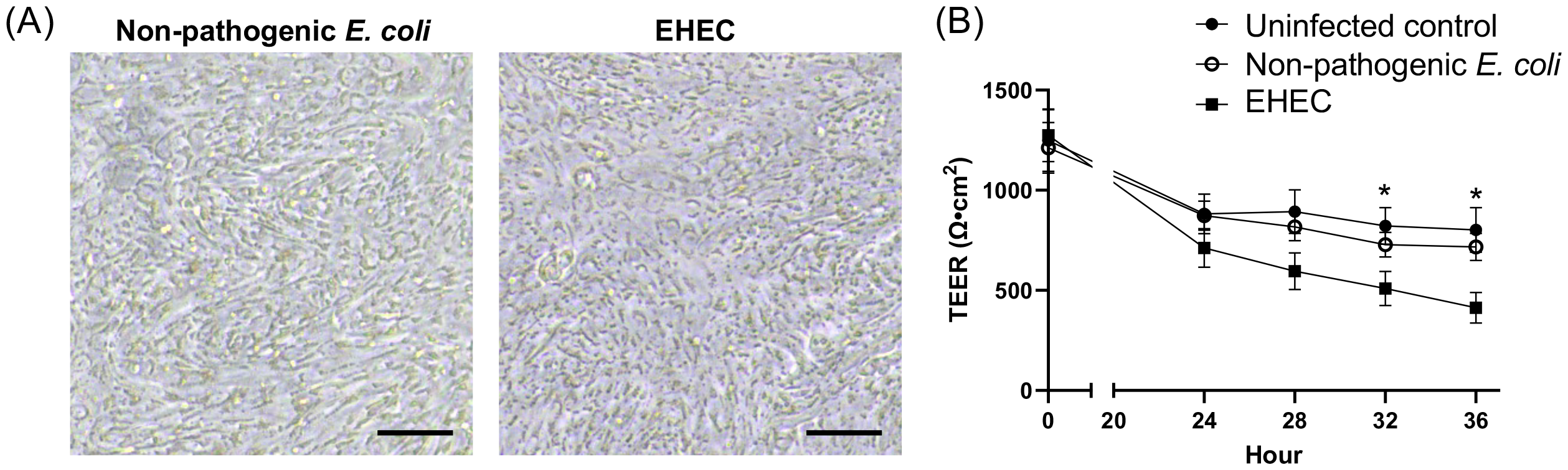
Evaluation of epithelial barrier integrity in canine colonoid-derived monolayers co-cultured with bacteria. (A) Representative phase-contrast image of colonoid-derived monolayers after 36 hours of co-culture with bacteria. Scale bar = 100 μm. (B) Changes in transepithelial electrical resistance (TEER) value since the initiation of co-culture. TEER values were measured up to 36 hours after the start of co-culture. Black circles indicate uninfected control group, white circles indicate non-pathogenic *E. coli* group, and black squares indicate EHEC group. * *P*<0.05.

### Expression of ZO-1, but not E-cadherin, decreased following EHEC co-culture

As previously reported, EHEC infection is known to downregulate tight junction protein expression and disrupt the gut barrier (15–17). We evaluated the expression levels of tight junction proteins, namely ZO-1 and E-cadherin, in canine colonoid-derived monolayers by using immunofluorescence staining after 36 hours of co-culture (Figure 4A). Fluorescence images of both non-pathogenic *E. coli* and EHEC cultures showed distinct tight junctions in the intercellular regions with no obvious disruption of tight junctions. To determine if there were differences in the intensity of tight junction expression, we performed line scans using fluorescence images to extract intercellular-specific fluorescence intensities and compared these intensities between the two groups (Figure 4B and 4C).

**Figure 4.**
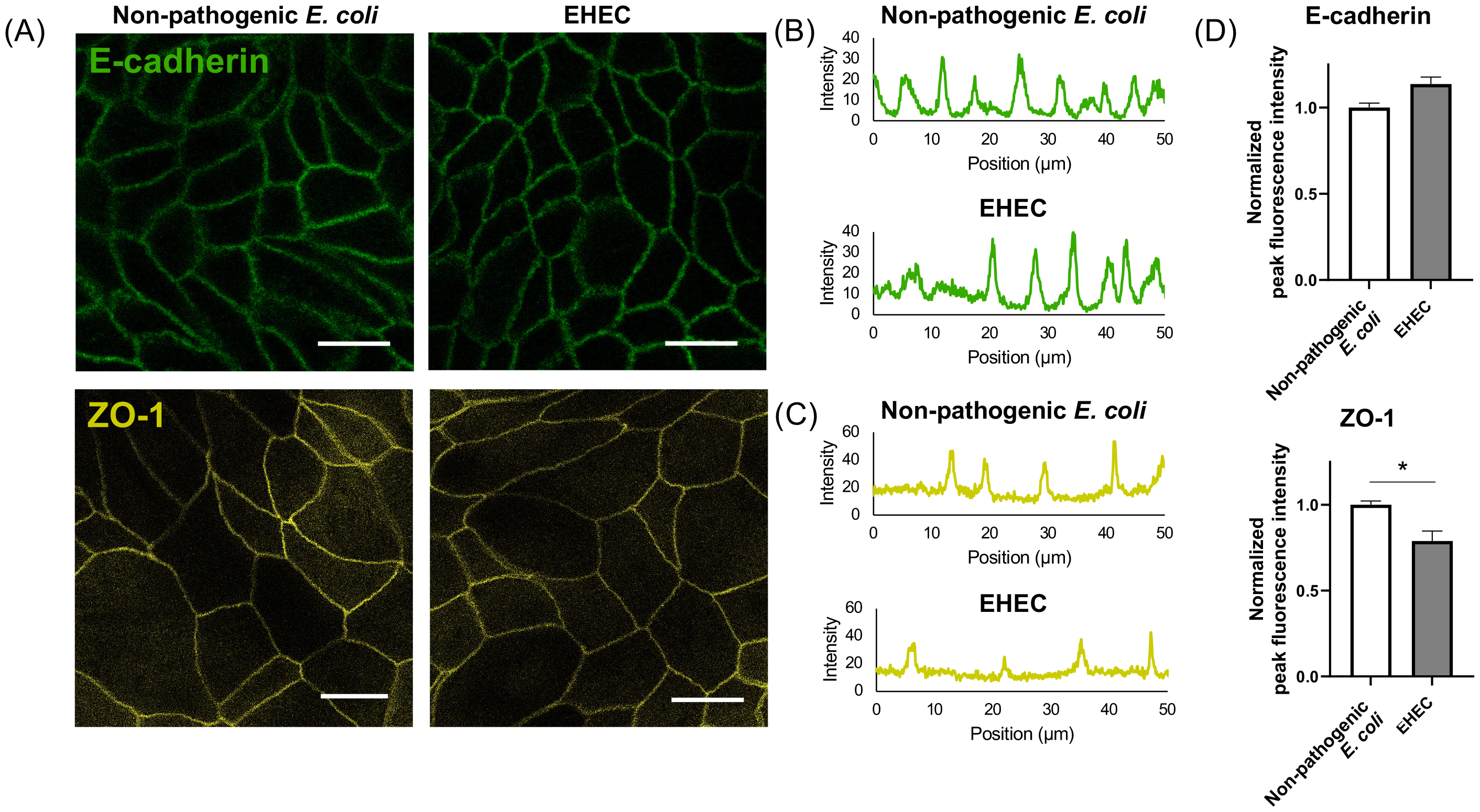
Tight junction proteins in canine colonoid-derived monolayers after bacterial co-culture. (A) Representative fluorescence image of tight junction proteins, E-cadherin (Green) and ZO-1 (Yellow), in colonoid-derived monolayers after 36 hours of co-culture with bacteria. Scale bar = 25 μm. (B) Representative line scan image of E-cadherin. The upper graph shows the Non-pathogenic *E. coli*group, and the lower graph shows the EHEC group. (C) Representative line scan image of ZO-1. The upper graph shows the Non-pathogenic *E. coli* group, and the lower graph shows the EHEC group. (D) Comparison of the peak intensity in the line scan. To evaluate intercellular specific fluorescence intensity, a comparison of the intensity of each peak on the line scan was performed. This measurement was performed using three biological replicates with three technical replicates. For all samples, images were obtained in five randomly selected fields of view, and a line scan was performed on each image. In each biological replicate, the fluorescence intensity was normalized by the mean of the peak fluorescence intensity in non-pathogenic *E. coli*. White bars indicate the non-pathogenic *E. coli* group, and gray bars indicate the EHEC group. The error bars represent the standard error of the mean. * *P*<0.05.

While the expression of E-cadherin was not different between the two groups (*P*=0.06), that of ZO-1 decreased significantly for EHEC-exposed monolayers (*P*=0.02) (Figure 4D).

### Mucus production in the canine colonoid-derived monolayers was not altered by EHEC infection

We assessed the mucus production of canine colonoid-derived monolayers after the co-culture with non-pathogenic *E. coli* or EHEC because changes in mucus-producing function have been reported in mice and humans (17, 18). After 36 hours of co-culture with the bacteria, SNA staining was performed and the number of SNA-positive cells per high magnification field of view was calculated. The number of SNA-positive cells per field of view was 2.8 ± 0.3 cells with the non-pathogenic *E. coli* co-culture and 2.4 ± 0.3 cells with the EHEC co-culture (*P*=0.27) (Figure 5).

**Figure 5.**
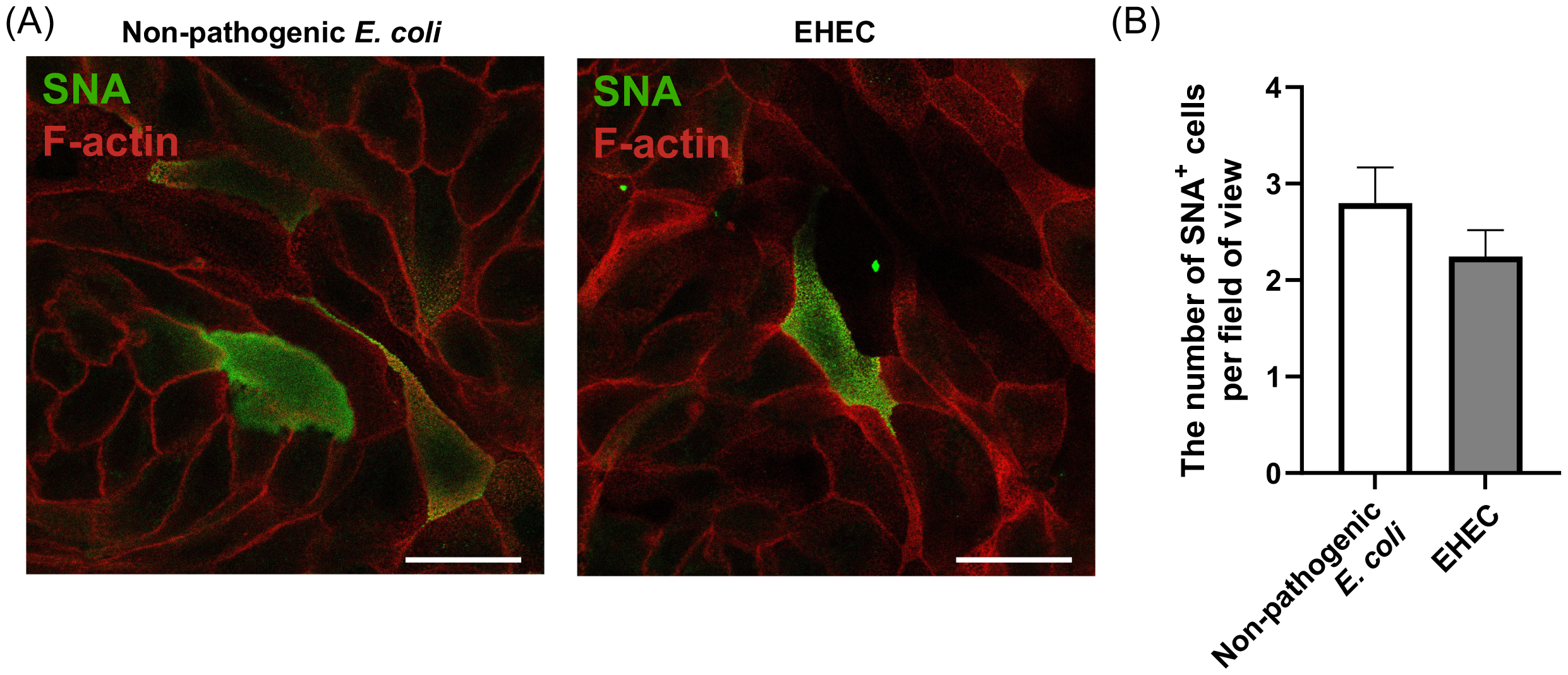
Evaluation of mucus production in canine colonoid-derived monolayers after bacterial co-culture. (A) Representative fluorescence image of SNA staining. SNA is a lectin that binds to sialic acid in the sugar chains of mucus, which could be interpreted to mean that SNA-positive cells are goblet cells. Scale bar = 25 μm. (B) Quantification of SNA-positive cells. To calculate the percentage of SNA-positive cells in each group, the number of SNA-positive cells per field of view under 63x magnification was counted. This quantification was performed in five randomly selected fields of view in each sample with three technical replicates using three biological replicates. White bars indicate the non-pathogenic *E. coli* group, and gray bars indicate the EHEC group. The error bars represent the standard error of the mean. * *P*<0.05.

### Inflammatory cytokine production was elicited by EHEC infection

We measured the concentration of IL-8 and TNF-α in the medium after 36 hours of co-culture with bacteria because these cytokines are upregulated in the intestinal tract after EHEC infection (19– 21). In accordance with previous studies, these measurements were performed by ELISA using basolateral medium (21). The concentration of IL-8 (782.7 ± 84.1 pg/ml) from exposure to non-pathogenic *E. coli* was similar to the concentration for the EHEC exposed monolayers (1,271.0 ± 448.9 pg/ml) (*P*=0.54). The concentration of TNF-α was significantly higher with EHEC co-culture (9.8 ± 1.6 pg/ml) compared with non-pathogenic *E. coli* co-culture (4.7 ± 1.4 pg/ml) (*P*=0.03, Figure 6).

**Figure 6.**
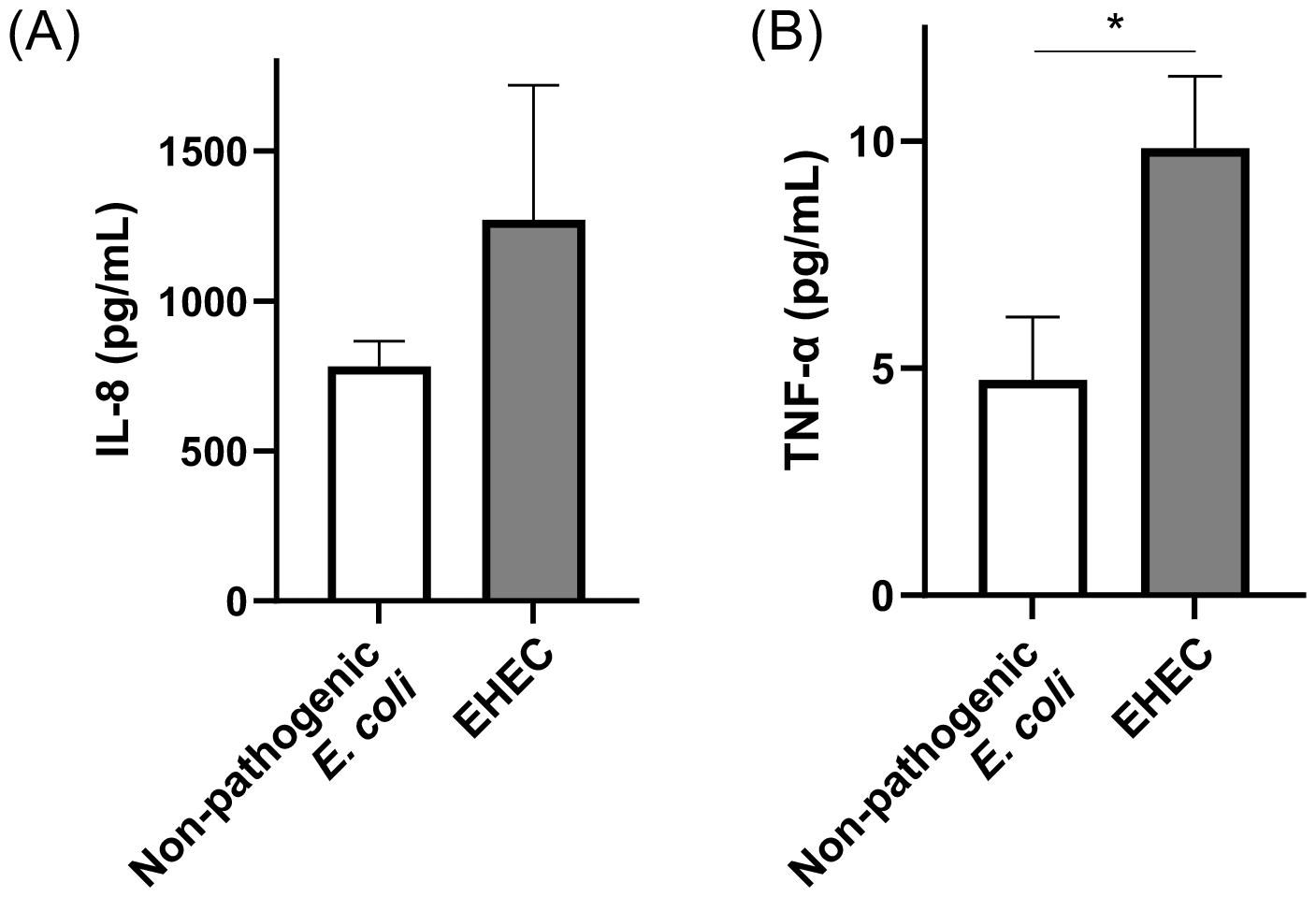
Inflammatory cytokine production in canine colonoid-derived monolayers co-cultured with bacteria. Concentration of inflammatory cytokine, IL-8 (A) and TNF-α (B) was measured using basolateral culture medium after 36 hours of co-culture by ELISA. White bars indicate the non-pathogenic *E. coli* group, and gray bars indicate the EHEC group. The error bars represent the standard error of the mean. * *P*<0.05.

## Discussion

This study introduces a novel canine colonoid-derived monolayer model for *in vitro* studies of EHEC infection. We have found that incorporating a saline-only incubation step effectively limits bacterial overgrowth without causing undesirable artefacts to the monolayers. Additionally, results from the non-pathogenic *E. coli* co-culture provides a useful contrast for evaluating specific effects of EHEC infection on canine intestinal epithelial cells.

Cattle are the primary reservoirs for EHEC and the transmission of this bacterium to humans primarily occurs through consumption of contaminated food products derived from cattle, the use of cattle manure as fertilizer, and water supplies contaminated by runoff from cattle farms (22). Additionally, dogs consuming raw-food diets are increasingly at risk of being contaminated with enteric pathogens, including antibiotic-resistant strains of *E. coli* and *Salmonella enterica* (23). In our study, we utilized an EHEC isolate sourced from cattle feces to more accurately mirror the natural infection route in dogs. This methodology provides a realistic representation of potential EHEC exposure in the canine environment.

In scanning electron microscopy (SEM) images, we observed the adherence of EHEC to epithelial monolayers generated from canine colonoids. However, the quantity of bacteria that adhered was relatively low. Previous research indicates that EHEC colonizes the epithelial surface by utilizing its flagella to attach to the mucus produced by intestinal goblet cells (24). This highlights the crucial role of intestinal mucus in EHEC infections. The monolayers in our study, however, were cultured in a stem cell maintenance medium, which resulted in a high population of stem cells and a markedly low population of goblet cells (25). This condition was anticipated to reduce the number of adhering bacteria.

Supporting this, a study involving human intestinal organoids demonstrated that EHEC adherence was significantly higher (by tenfold) in monolayers with increased mucus secretion, achieved through induced differentiation, compared to those cultured in a stem cell maintenance medium (26). This suggests the importance of developing methods to differentiate canine organoid-derived monolayers for more effective studies on EHEC infection in the future.

The Centers for Disease Control and Prevention (CDC) estimates that foodborne infections caused by EHEC result in approximately 63,000 cases annually in the United States, leading to over 2,100 hospitalizations and fatalities (27). The economic impact of these illnesses, considering medical costs, loss of life, and reduced productivity, is estimated at $405 million each year (28). Focusing specifically on severe EHEC infections, about 10% of patients with infections from Shiga toxin-producing *E. coli* (STEC) may develop HUS, with a mortality rate of 3 to 5% (29). In canine subjects, severe EHEC infections leading to HUS have been documented, though predominantly in experimental settings or dogs with compromised immunity (4, 5). In our study, the impact of EHEC infection included a modest decrease in TEER, with a reduction of approximately 500 Ω*cm2. This observation was consistent with other relatively mild alterations that we observed, aligning with the typical progression of EHEC infections in dogs under natural conditions.

Various *in vitro* studies have reported a reduction in colon mucus in cell culture (25, 26), yet *in vivo* information remains scarce. Research has shown that EHEC can gain a competitive edge by utilizing mucus-derived sugars, serving not only as a carbon source but also as signaling metabolites within the intestinal microenvironment (30). This enables EHEC to modify its proliferative and virulent behavior in response to mucus presence (31). However, there is a lack of data regarding mucus changes in dogs with EHEC infection. Thus, our observations, which found no discernible differences in mucus between infections with non-pathogenic *E. coli* and EHEC, may not be entirely unexpected. Further research is required to understand mucus dynamics in canine EHEC infection and its role in the pathogenesis of the disease.

In children infected with EHEC, there is a correlation between increased levels of the proinflammatory cytokine IL-8 in the blood and a heightened risk of developing HUS (32). However, this relationship has not been extensively studied in dogs with EHEC infections. In our research, we did not observe a significant rise in IL-8 levels in canine colonoid-derived monolayers infected with EHEC. This absence of a marked increase in IL-8 suggests that the response to EHEC infections in dogs may mirror the generally milder course of EHEC infections seen in dogs under natural conditions. A notable model for studying EHEC infections in humans is the murine model of *Citrobacter rodentium* infection (33, 34). Similar to what we observed in our study, TNF-α levels increase in the gut for this model. Since TNF-α is a known proinflammatory cytokine that regulates macrophage function and these macrophages are protective factors against bacterial infection (35), it seems reasonable that TNF-α expression is induced by EHEC infection *in vivo* in mice and *in vitro* in dogs. However, the expression of TNF-α in mice *in vivo* and in dogs *in vitro* has not been shown. The interaction with immune cells cannot be discussed in this co-culture model of epithelial cells and bacteria alone, so more complex culture systems, such as Gut-on-a-Chip, are needed.

Recent research has explored species-specific variations in EHEC pathogenesis between mice and humans using Gut-on-a-Chip microfluidic technology with human colonoids. This study identified specific microbiome metabolites that contribute to the species-specific susceptibility to EHEC (36). Building on this, we have successfully established a canine Gut-on-a-Chip model using canine intestinal organoids (10). Should we observe parallels in the canine model, especially when comparing intestinal microbiomes from dogs, humans, and potentially mice, this approach could significantly enhance our understanding of microbiome-derived metabolites in EHEC pathogenesis.

In conclusion, our study marks a critical step forward in deciphering the complexities of EHEC pathogenesis, with a special focus on canine models. The introduction of a novel canine colonoid-derived monolayer model lays the groundwork for a more nuanced comparative analysis of EHEC strains during infections. Our approach, which includes the strategic modification of culture conditions by using saline-only media, successfully limits bacterial overgrowth that typically compromises *in vitro* co-culture experiments. This model, along with a novel canine Gut-on-a-Chip model (10), represent an important opportunity to better understand the role of microbiome metabolites during EHEC infection across different species. The insights gained from such investigations have the potential to guide more effective strategies for the prevention and treatment of EHEC infections, thereby benefiting both animal and human health.

## Materials and Methods

### Animals

For this study, three clinically healthy dogs undergoing dental procedures at the Washington State University (WSU) Veterinary Teaching Hospital (VTH) Community Practice Service were included. These dogs, aged between 1 and 12 years, were selected based on comprehensive physical examinations, blood work, and no history of chronic diseases affecting the heart, kidneys, liver, or intestines. Only those dogs deemed fit for elective procedure under general anesthesia were included in the study. This study was conducted with the approval of the Washington State University Institutional Animal Care and Use Committee (IACUC Approval: ASAF#6993), with details on the dogs provided in Supplementary Table 1.

### Generation of canine colonoids

Colonic stem cells were isolated from the biopsied colonic tissue using the previously reported protocol (8). Briefly, biopsied colonic tissues were cut into small pieces and treated with 30 mM EDTA solution (Invitrogen) for 60 min at 4°C to isolate crypts containing stem cells. These crypts were collected, washed using ice-cold Dulbecco’s phosphate-buffered saline (PBS, Gibco) with 1x penicillin/streptomycin (Gibco), embedded in Matrigel (Corning), and then seeded into 48-well plates (Thermo Scientific). After the Matrigel domes were solidified in a 37°C incubator, 300 μl of organoid culture medium were applied to each well. For composition of organoid culture medium, we used DMEM/F12 (Gibco) supplemented with 2 mM GlutaMAX (Gibco), 10 mM HEPES (Gibco), 1x penicillin/streptomycin (Gibco), 10% (vol/vol) conditioned medium of Noggin (37), 20% (vol/vol) conditioned medium of R-spondin, 100 ng/ml recombinant murine Wnt-3a (PeproTech), 50 ng/ml murine Epidermal Growth Factor (EGF) (PeproTech), 10 nM gastrin (Sigma-Aldrich), 500 nM A-83-01 (Sigma-Aldrich), 10 μM SB202190 (Sigma-Aldrich), 1 mM N-acetyl-L-cysteine (MP Biomedicals), 10 mM nicotinamide (Sigma-Aldrich), 1x B27 supplement (Gibco), 1x N2 MAX media supplement (R&D Systems), and 100 μg/ml Primocin (Invitrogen). For the initial two days following crypt isolation, 10 μM Y-27632 (Stem Cell Technologies) and 2.5 μM CHIR 99021 (Stem Cell Technologies) were added to the culture medium. Once the colonoids maturated, the Matrigel dome was dissolved using Cell Recovery Solution (Corning), and the colonoids were collected and treated with TrypLE Express (Gibco) and reseeded into 48-well plates.

### Development of canine colonoid-derived monolayers

Once the colonoids reached maturity, the Matrigel dome was dissolved using Cell Recovery Solution (Corning) and colonoids were collected by centrifugation (200x *g*, 5 min, 4°C). Collected colonoids were treated with TrypLE Express containing 10 μM of Y-27632 for 10 min at 37 °C. Subsequently, the dissociated colonoids were filtered through a 70 μm cell strainer (Fisher Scientific) to obtain single cells and the cells were centrifuged at 200x *g*, 4°C for 5 min.

For the preparation of cell culture inserts (Falcon), these inserts were coated using 100 μg/ml Matrigel and 30 mg/ml collagen I (Gibco) in DMEM/F12 supplemented with 2 mM GlutaMAX, 10 mM HEPES, and 1x penicillin/streptomycin. Dissociated single cells obtained from colonoids were suspended with organoid culture medium supplemented with 10 μM Y-27632 and 2.5 μM CHIR 99021 and seeded onto the pre-coated cell culture insert at a concentration of 1x10^6^ cells/ml. The monolayers were cultured in organoid culture medium containing 10 μM Y-27632 and 2.5 μM CHIR 99021 until the day after seeding, after which organoid culture medium was changed every other day.

### Microbial culture for infection experiment

This study used an EHEC isolate from cattle to challenge the monolayer and a non-pathogenic *E. coli* strain, MC4100, as an infection control. MC4100 is a derivative of strain K-12, that was originally isolated from a human in 1922 (14) The EHEC isolate was PCR positive for *eae, h7, stx1*, and *stx2/vt2* (Supple Figure 2) (38). Both strains of bacteria were cultured overnight in sterile Luria-Bertani (LB) medium at 37°C under shaking at 200 rpm. Subsequently, 100 μl of the overnight culture was transferred to 3-ml fresh LB medium and incubated for 3 h to bring the bacteria to the logarithmic growth phase. After incubation, the bacterial solution was centrifuged at 10,000x *g* for 3 min, washed with normal saline (0.9% W/V NaCl), and further centrifuged, and resuspended to 2.0 x 10^7^ CFU/ml in saline.

### Co-culture of bacteria and canine colonoid-derived monolayers

Four days after the start of culture of the colonoid-derived monolayer, the culture media on both the apical and basolateral chambers were removed and washed three times with saline (Figure 1A). Saline (200 μl) was added to the apical chamber, and 500 μl of organoid culture medium depleted of antibiotics was added to the basolateral chamber. Then, the colonoid-derived monolayers were further incubated (37°C, 24 h). After incubation, the bacterial suspension was added to apical chamber to a final concentration of 1.0 x 10^6^ CFU/ml.

### Assessment of intestinal barrier integrity

The intestinal barrier integrity of the colonoid-derived monolayers was evaluated by measuring the TEER value following exposure to bacteria. Electrical resistance (Ω_t_) was quantified using Ag/AgCl electrodes connected to a Volt-Ohm meter (Millicell ERS-2, Millipore), and this value was then transformed into a TEER measurement using the following equation: TEER = (Ω_t_ - Ω_blank_) x A, where Ω_blank_ represents the resistance of the blank well in ohms, and A denotes the surface area of the culture insert in cm^2^.

### Immunocytochemistry of canine colonoid-derived monolayers

Colonoid-derived monolayers were fixed after 36 h co-culture by using 4 % PFA (Thermo Scientific), followed by membrane permeabilization with 0.3 % Triton-X (Thermo Scientific), blocking with 2 % bovine serum albumin (Cytiva), and then addition of primary antibodies. For visualization of E-cadherin and ZO-1, monoclonal anti-mouse E-cadherin antibody (36/E-cadherin, BD Biosciences) and polyclonal anti-rabbit ZO-1 antibody (61-7300, Invitrogen) were used. SNA staining (Vector laboratories) was used to visualize goblet cells. The colonoid-derived monolayers were then washed with PBS and secondary antibodies were added (Anti-Rabbit IgG H&L labeled with Alexa Fluor 555). After further washing, the nuclei and F-actin were visualized using DAPI and Alexa Fluor 647 Phalloidin (Thermo Fisher Scientific), respectively. Then, the monolayer was mounted by Prolong Gold Antifade reagent (Thermo Fisher Scientific) and imaged using a white-light point scanning confocal microscope (SP8-X, Leica). Fluorescence images were acquired under 63× objective, with excitation laser sources of 405 nm, 499 nm, 553 nm, and 653 nm and high efficiency Leica HyD detector, and processed using LAS X software (Leica). Intensity of fluorescence signal for E-cadherin and ZO-1 was quantified by performing line scan using ImageJ 1.54d (39). Mean of maximum intensity value of each peak was determined by taking the peak intensity value from at least four peaks per line scan and normalizing it by the value obtained from non-pathogenic *E. coli* infected monolayers. The number of SNA-positive cells per field of view was used to evaluate the number of goblet cells. These evaluations were performed on five randomly selected fields of view from each of the three biological replicates, each of which consisted of three technical replicates.

### Scanning electron microscopy

The colonoid-derived monolayers were fixed with 2.5 % (vol/vol) glutaraldehyde (Ted Pella) in 0.1 M sodium cacodylate buffer (Ted Pella) overnight at 4°C. The samples were then rinsed with 0.1 M cacodylate buffer and fixed with 1 % osmium tetroxide (Electron Microscopy Sciences) in 0.1 M sodium cacodylate buffer for 30 min at room temperature, followed by serial dehydration in 30-100 % ethanol and hexamethyldisilazane (HDMS) (SPI Supplies). Subsequently, the samples were mounted on stubs, coated with Pt/Pd using a high-resolution sputter coater (Cressington), and imaged using Quanta 200F SEM (FEI).

### Enzyme-linked immunosorbent assay

After the infection experiment was completed, apical and basolateral media were collected. After centrifugation to remove the cellular components of the medium, both media were frozen at -80°C until the inflammatory cytokines were determined by ELISA. The IL-8 Canine ELISA Kit (#ECCXCL8, Invitrogen) and TNF-α Canine ELISA Kit (# ECTNF, Invitrogen) were used to measure IL-8 and TNF-α, respectively, and the assays were performed according to the manufacturer’s protocol. In accordance with previous studies, these measurements were made using basolateral medium (21).

### Statistical analysis

The study was tested in three independent technical replicates using three biological replicates in all analyses. Statistical analysis was conducted using R v4.1.0 (R core team) and figures were generated using GraphPad Prism 10.0.2.232 (Dotmatics). To confirm the normality of each dataset, the Shapiro-Wilk’s test was performed. The Kruskal-Wallis and Dunnett tests were employed for the comparison of TEER value among each condition. The Wilcoxon test was utilized to compare the concentration of IL-8. Additionally, Student t-tests were applied to compare the number of SNA-positive cells, fluorescence intensity of tight junction proteins, and the concentration of TNF-α. All results were presented as mean ± standard error of the mean (SEM). A *P*<0.05 was considered statistically significant.

## Supplemental Material

**Supplementary Figure 1. Gene expression of differentiated cell markers in colonoid-derived monolayers**.

After three days of saline incubation, the expression levels of differentiated cell marker genes (*leucine-rich repeat containing G protein-coupled receptor 5* (*LGR5*), *Chromogranin A* (*CgA*), and *Lysozyme*) in the colonoid-derived monolayers were quantified by RT-qPCR. Complete medium (CM), a stem cell maintenance medium, was used for comparison. The error bars represent the standard error of the mean.

**Supplementary Figure 2. Characterization of enterohemorrhagic *E. coli***.

PCR results for toxin-related genes in enterohemorrhagic *E. coli* used in this study are shown. Animal 1 is the bovine-derived enterohemorrhagic *E. coli* used in this study, and Animals 2-4 are *E. coli* isolates that were collected from other animals but that tested negative. Known enterohemorrhagic *E. coli* were used as positive control (PC) 1 and 2, and a band of toxin-related genes of the same size as the positive control was identified in Animal 1.

## Acknowledgments

Authors would like to thank Small Animal Internal Medicine (Dr. Jillian Haines, Dr. Sarah Guess, Shelley Ensign LVT, Sybil Fiedler VTA) and Community Practice (Dr. Jessica Bell, Dr. Cassidy Cordon, Dr. Matt Mason, Mrs. Melody Gerber, Ms. Maggie deSouza, Ms. Becky Brodie) services at WSU VTH and WSU VTH Clinical Studies Coordinator Valorie Wiss for their support in case recruitment and sample collection from citizen scientists (patient donors). Authors would also like to thank Dr. Craig S. McConnel (Field Disease Investigation Unit at WSU) for providing EHEC bacteria isolated from bovine, Jenn Horton and Lindsay Parrish for their technical help in bacterial preparations and logistical support, and Dr.

Valerie Lynch-Holm and Dr. Brittney Wager from the Franceschi Microscopy and Imaging Center at WSU for their technical support in electron microscopy. This work was supported in part by the Office of The Director, National Institutes Of Health (K01OD030515 and R21OD031903 to Y.M.A.) and Japan Society for the Promotion of Science Overseas Challenge Program for Young Researchers (202280196 to I.N.).

